# Mutation of a conserved Gln residue does not abolish desensitization of acid-sensing ion channel 1

**DOI:** 10.1101/2020.12.19.423606

**Authors:** Matthew L Rook, Megan Miaro, Tyler Couch, Dana L Kneisley, Maria Musgaard, David M. MacLean

## Abstract

Desensitization is a common feature of ligand-gated ion channels although the molecular cause varies widely between channel types. Mutations that substantially reduce or abolish desensitization have been described for many ligand-gated ion channels including glutamate, GABA, glycine and nicotinic receptors but not for acid-sensing ion channels (ASICs) until recently. Mutating Gln276 to a glycine in human ASIC1a was reported to mostly abolish desensitization at both the macroscopic and single channel levels, potentially providing a valuable tool for subsequent studies. However, we find that in both human and chicken ASIC1 the effect of Q276G is modest. In chicken ASIC1, the equivalent Q277G slightly reduces desensitization when using pH 6.5 as a stimulus but desensitizes essentially like wild type when using more acidic pH values. In addition, steady-state desensitization is intact, albeit right-shifted, and recovery from desensitization is accelerated. Molecular dynamics simulations indicate that the Gln277 side chain participates in a hydrogen bond network that might stabilize the desensitized conformation. Consistent with this, destabilizing this network with the Q277N or Q277L mutations largely mimics the Q277G phenotype. In human ASIC1a, Q276G does not substantially reduce desensitization but surprisingly slows entry to and exit from the desensitized state, thus requiring longer agonist applications to reach equilibrium. Our data reveal that while the Q/G mutation does not substantially impair desensitization as previously reported, it does point to unexpected differences between chicken and human ASICs and the need for careful scrutiny before using this mutation in future studies.

## Introduction

Desensitization is a near-ubiquitous feature of ligand-gated ion channels (LGICs), which was first described more than 60 years ago^1^. In general, desensitization is thought to act as a protective mechanism, terminating aberrant signaling although other roles are possible^2–4^. As such, the molecular basis of desensitization has been a subject of inquiry for every type of LGIC. Mutations that essentially abolish or substantially reduce desensitization have been reported for glutamate, GABA, glycine and nicotinic receptors^5–10^. While there have been controversies surrounding the microscopic mechanisms of particular cases^11^, these mutations have been enormously helpful in driving structure-function investigations of desensitization as well as the insight into the physiological role^12^. Until recently, no such mutations had been reported for acid-sensing ion channels (ASICs).

ASICs are sodium-selective pH-activated trimeric ion channels. They are expressed widely in the central and peripheral nervous systems, as well as other tissues^13^. Given the ubiquity of the ligand, it is unsurprising that ASICs are implicated in a host of physiological processes and disease states including ischemic cell death, fear and anxiety, learning and memory, pain, muscle fatigue, migraine, bone morphogenesis, inflammation and cancer^14–16^. In mammals, the ASIC family includes four proton-sensitive members: ASIC1a, ASIC1b, ASIC2a and ASIC3. The individual subunits all have the same topology with intracellular amino and carboxy terminal tails of approximately 20 to 80 amino acid residues, separated by a large extracellular domain, two transmembrane helices and a small amino terminal re-entrant loop^17,18^. The extracellular domain is divided into distinct thumb, finger, knuckle, palm, and β-ball domains (Figure 1A). ASIC activation by acidic conditions is believed to occur through protonation of distinct residues in the interface between the thumb and finger as well as a cluster of acidic side chains in the palm domain^17,19–21^. Protonation also triggers desensitization, either with mild acidic stimuli (pH in the 7.4-6.9 range), which leads to steady-state desensitization (SSD) in the absence of channel activation, or with strong stimuli (i.e., pH 6.8-4) which also opens the channel. Desensitization depends on the isomerization or swivel of a critical linker in the palm domain, which connects the 11^th^ and 12^th^ β strands^22,23^. This linker is composed of Leu414 and Asn415 (Figure 1B) and in the resting and open states, the Leu residue points outward, away from the central axis of the channel. However, in the desensitized state these amino acid residues essentially switch positions, with Leu414 swiveling downward and in towards the central axis. It has been suggested that Gln276 (Gln277 in chicken ASIC1) acts as a valve to prevent linker rotation, stabilizing the desensitized state and, furthermore, that eliminating the 276 side chain using the Q276G mutation in effect produces a ‘leaky’ valve which enables channels to readily escape desensitization and remain open^24^.

**Figure 1.**
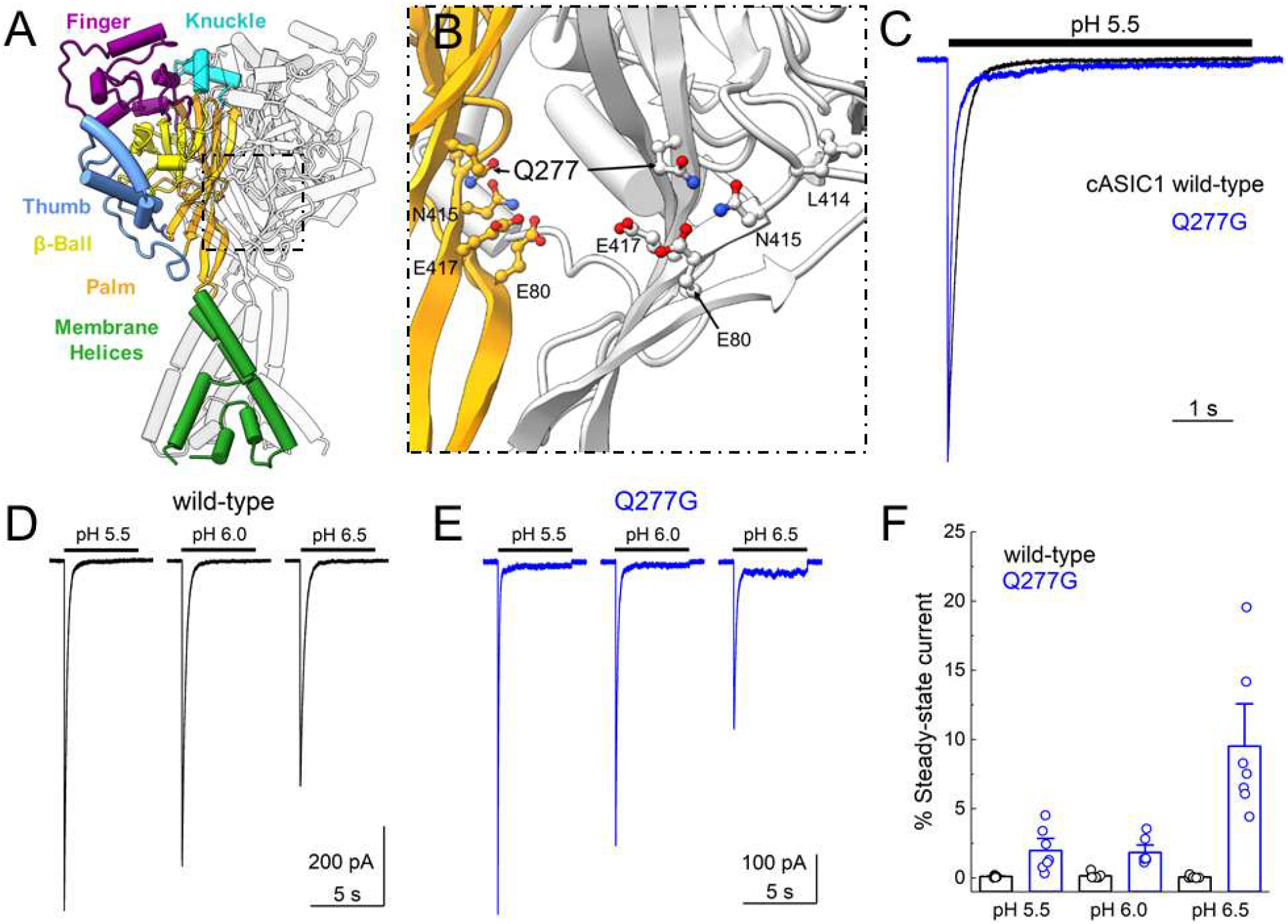
cASIC1 Q277G exhibits strong desensitization over several pH values. **(A)** Structure of the cASIC1 resting state (PDB: 6VTL). Domains are identified by color in one subunit while the remaining two subunits are colored light or darker grey. **(B)** Close in view of the boxed region in **(A)** showing Q277 position in two subunits as well as functionally relevant amino acids. The ‘front’ subunit has been removed leaving only the colored and ‘rear’ subunits for clarity. **(C)** Peak normalized outside-out patch responses from cASIC1 wild type (*black trace*) or Q277G (*blue trace*) during a jump from pH 8 to pH 5.5. **(D & E)** Responses from single outside-out cASIC1 wild type **(D)** or Q277G **(E)** patches to the indicated pH stimuli. **(F)** Summary of the percent steady-state current, normalized to the peak response within a pH, over several patches. Circles denote individual patches and error bars show S.E.M.

To further study the Q276G mutant and relate it to the majority of structural data, we tested the Q276G equivalent in cASIC1 (Q277G) using piezo-driven fast perfusion in excised patches. We found that when using pH 6.5 to open the channels, cASIC1 Q277G does have slightly reduced desensitization, however, when using more acidic stimuli Q276G behaves essentially like wild type with desensitization principally intact. Moreover, we found that Q277G accelerates recovery from desensitization by orders of magnitude and reduces the apparent stability of the desensitized state. Based on molecular dynamics simulations, we hypothesize that Gln277 coordinates a series of hydrogen bonds within the palm domain, thereby stabilizing the desensitized conformation. Consistent with this electrostatic mechanism, a slight mutation to Q277N or Q277L also accelerates recovery from desensitization. Finally, we find that hASIC1a Q276G exhibits robust pH-dependent desensitization, in contrast to prior work.

## Materials and Methods

### HEK ASIC knockout cell creation

A guide RNA sequence (GGCTAAAGCGGAACTCGTTG-PAM) targeting the coding region of *ASIC1* was cloned into Bbsl-linearized pSpCas9(BB)-2A-GFP vector (a kind gift from Feng Zhang, Addgene plasmid #48138) as previously described^25^. Transfected human embryonic kidney cells (HEK293, ATCC number CRL-3216) cells were verified for GFP expression and clonally expanded following serial dilution. Clonal lines were screened for on-target genome editing by Sanger sequencing of PCR products (Fwd TTGGAGGAACCCTGGATGTGTC, Rev TAACTCCTCTGCTGTGAGTGGC). Knock-out was confirmed by Western blotting. Briefly, 10^5^ cell equivalents of RIPA lysate from the parental cell line, knockout clone, and clone transiently transfected with human ASIC1a cDNA were resolved on an acrylamide gel and transferred to nitrocellulose membranes. Blots were blocked with bovine serum albumin and probed with an ASIC1-specific antibody (NeuroMab clone N271/44) overnight at a 1:1000 dilution. Blots were washed with Tris-buffered saline supplemented with 0.1% Tween-20 and probed with goat anti-mouse IgG HRP-conjugated secondary. Blots were imaged with an Azure 300 Imaging System. After inactivation of HRP with sodium azide, the blot was probed again with Direct-Blot HRP anti-GAPDH (Biolegend) antibody as a loading control.

### Cell culture, mutagenesis and transfection

HEK293T ASIC knockout (KO) cells were maintained in Dulbecco’s Modification of Eagle’s Medium (DMEM) with 4.5 g/L glucose, L-glutamine & sodium pyruvate (Corning/Mediatech, Inc.) or Minimum Essential Medium (MEM) with Glutamax & Earle’s Salts (Gibco), supplemented with 10% FBS (Atlas Biologicals) and penicillin/streptomycin (Invitrogen). Cells were passaged every 2 to 3 days when approximately 90% confluence was achieved. HEK293 KO cells were plated on tissue culture treated 35 mm dishes, transfected 24 to 48 hours later and recorded from 12-48 hours post-transfection. Cells were transiently transfected with the indicated ASIC construct and eGFP using an ASIC:eGFP ratio of between 2.5 - 10:1 μg of cDNA per 10 mL of media, depending on the construct. For hASIC1a wild type whole cell experiments (Figure 6), a ratio of 0.25:0.25:1 ug of hASCI1a, eGFP and pUC empty vector was used. Transfections were performed using polyethylenimine 25k (PEI 25k, Polysciences, Inc) following manufacturer’s instructions, with media change at 1 to 8 hours post-transfection. Mutations were introduced using site-directed mutagenesis PCR and confirmed by sequencing (Fisher Scientific/Eurofins Genomics).

### Electrophysiology

Culture dishes were visualized with phase contrast on a Nikon Ti2 microscope using a 20x objective. GFP was excited using a 455 nm or 470 nm LED (Thorlabs) and dichroic filter cube for emission detection. Outside-out patches were excised using heat-polished, thick-walled borosilicate glass pipettes of 3 to 15 MΩ resistance. The pipette internal solution contained (in mM) 135 CsF, 33 CsOH, 11 EGTA, 10 HEPES, 2 MgCl_2_ and 1 CaCl_2_ (pH 7.4). External solutions with pH values greater than 7 were composed of (in mM) 150 NaCl, 10 HEPES, 1 CaCl_2_ and 1 MgCl_2_ with pH values adjusted to their respective values using NaOH. For solutions with a pH value lower than 7, HEPES was replaced with MES. All recordings were performed at room temperature with a holding potential of −60 mV using an Axopatch 200B amplifier (Molecular Devices). Data were acquired using AxoGraph software (Axograph) at 20 kHz, filtered at 10 kHz and digitized using a USB-6343 DAQ (National Instruments). Series resistance was routinely compensated by 90 to 95% where the peak amplitude exceeded 100 pA. Rapid perfusion was performed using home-built, double- or triple-barrel application pipettes (Vitrocom), manufactured according to a prior method^26^. Application pipettes were translated using piezo actuators driven by voltage power supplies. The command voltages were generally low pass filtered (50-100 Hz, eight-pole Bessel). Whole cell recording (Supplemental Figure 1, Figure 6) used identical conditions except patch pipette and application pipette diameters tended to be larger.

### Molecular Dynamics Simulations

The systems for molecular dynamics simulations were constructed using the cASIC1 structure proposed to illustrate the desensitized state (PDB ID: 4NYK; resolution: 3.00 Å)^27^. Residues 42-455 were resolved in the crystal structure; of these 23 residues had missing atoms which were added using Modeller v9.21^28^. The model was oriented for placement in a lipid bilayer by aligning the complete structure with the corresponding structure from the Orientations of Proteins in Membranes (OPM) database^29^. Protonation states of specific residues were set using *pdb2gmx* during the system setup in GROMACS^30^. In all cases, residues Glu98, His111, Glu239, His328, Glu354, Glu374, Asp408 and Asp433 were protonated, leaving the acidic residues neutral and histidine residues with a positive charge. This was considered the “background” protonation setup and the purpose was to maintain an overall stable protein structure. However, the importance of the presence of these individual protons was not tested in this study as they are relatively far away from our region of interest. On the given background, Glu80, Glu412 and Glu417 were protonated or deprotonated in accordance with the table below to test the importance of protonation of these specific residues.

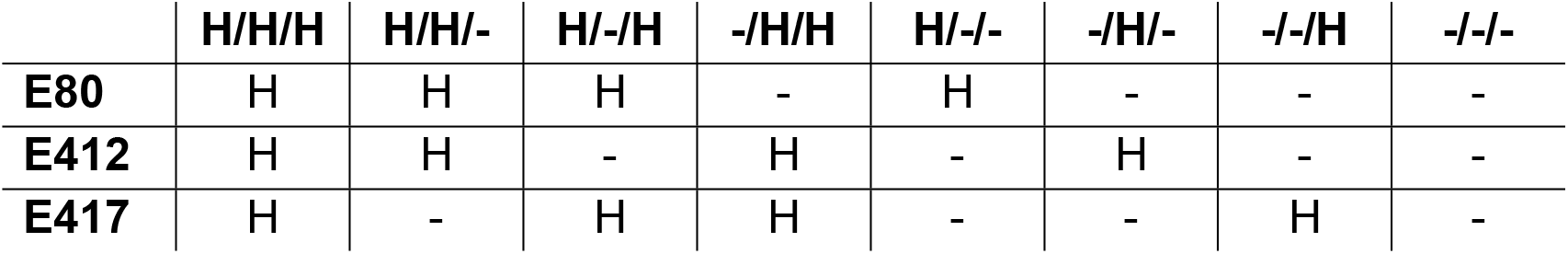

The Charmm36m force field was applied^31^. The initial POPC lipid bilayer (120 Å × 120 Å) was generated using the membrane builder of the CHARMM-GUI with 4NYK inserted using the replacement method^32^. The protein structures with different protonation states were then inserted into this original lipid bilayer using the InflateGro method^33^. The crystallographic water molecules and chloride ions were retained. Water molecules (TIP3P model^34^) were further generated to fill the box (120 Å × 120 Å × 161 Å) with solvent, and sodium and chloride ions were added to neutralize the system at a concentration of 0.15 M NaCl.

The simulations were performed using GROMACS 2019.4^30^. All systems were minimized until convergence or to a maximum of 5000 steps. The systems were then equilibrated in six steps totaling to 375 ps, using the standard method from the CHARMM-GUI. The first three equilibration runs used a time step of 1 fs while the last three and the production run used a time step of 2 fs. The first three equilibration runs were each 25 ps long and the final three were each 100 ps long. The position restraints were gradually lifted during the equilibration steps as suggested in the default CHARMM-GUI protocol. Periodic boundary conditions were applied. The Verlet cutoff scheme was used throughout with a force-switch modifier starting at 10 Å and a cutoff of 12 Å. A cutoff of 12 Å was used for short-range electrostatics and the particle mesh Ewald (PME) method was used for long-range electrostatics^35,36^. A Berendsen thermostat was used for all steps of the equilibration and a Nose-Hoover thermostat^37,38^ was utilized in the production run to maintain the temperature at 310.15 K for all steps. Using semi-isotropic pressure coupling, the pressure was maintained at 1 bar in the last four steps of equilibration and in the production run using the Berendsen barostat^39^ and the Parrinello-Rahman barostat^40^, respectively. The LINCS algorithm was used to constrain covalent bonds to hydrogen atoms^41^. The production runs were 100 ns long with a total of three repeats for each system. Each repeat had different initial velocities.

The system for the Q277N mutant was prepared as above, with the exception that the Gln277 sidechain was manually mutated to Asn by prior to system setup. The same background protonation scheme was used, and additionally Glu412 and Glu417 were protonated. The system was simulated as described with three repeats of 100 ns each.

Potential hydrogen bonds between the residues Glu80, Gln277, Glu412, Leu414 and Glu417 were sampled every 10 ps. The donor atoms include: Q277NH1, Q277NH2, E80HE2 (if protonated), E412HE2 (if protonated), and E417HE2 (if protonated). The acceptor atoms include: Q277OE1, E80OE1, E80OE2, E412OE1, E412OE2, L414O, E417OE1 and E417OE2. The 4.3.1 Hydrogen Bond Analysis module^42^ of MDAnalysis^43,44^ was used for the analysis, employing an updated and adapted version of M. Chavent’s Jupyter Notebook available on GitHub (https://github.com/MChavent/Hbond-analysis)^45^. Default cutoffs were used for the donor-acceptor distance (3.0 Å) and the donor-hydrogen-acceptor angle (150°). The presence of each unique hydrogen bond was calculated over the trajectory and expressed as a percentage of the total trajectory; the presence of equivalent hydrogen bonds (e.g., from 417OE1 and 417OE2 in the deprotonated state) were added to give one overall percentage for the given interaction. Plots were prepared using the Matplotlib package in Python. Figures were prepared using VMD^46^.

### Statistics and Data Analysis

Current desensitization decays were fitted using exponential decay functions in Clampfit (Molecular Devices). The percent of steady-state current was the current at the end of a pH application which had reached equilibrium divided by the peak current. For recovery from desensitization experiments, the test peak (i.e., the second response) was normalized to the conditioning peak (i.e., the first response). OriginLab (OriginLab Corp) was used to fit the normalized responses to:

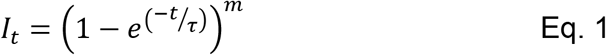

Where *I*_*t*_ is the fraction of the test peak at an interpulse interval of *t* compared to the conditioning peak, *τ* is the time constant of recovery and *m* is the slope of the recovery curve. Each protocol was performed between 1 and 3 times on a single patch, with the resulting test peak/conditioning peak ratios averaged together. Patches were individually fit and averages for the fits were reported in the text. N was taken to be a single patch. For activation and steady-state desensitization curves (SSD), peak currents within a patch were normalized to the peak response evoked by pH 5.5 and fit to:

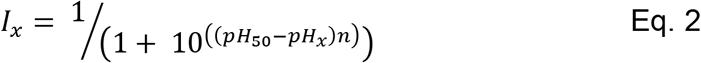

where *I*_*x*_ is the current at a given pH value X, *pH*_*50*_ is the pH yielding half maximal response and *n* is the Hill slope. Patches were individually fit and averages for the fits were reported in the text. N was taken to be a single patch.

Unless otherwise noted, statistical testing was done using nonparametric permutation or randomization tests with at least 100,000 iterations implemented in Python to assess statistical significance. Statistical comparisons of recovery from desensitization were based and reported on differences in recovery time constant.

## Results

Our goal was to investigate the functional properties of the Q276G mutation in a cASIC1 background, to permit easy comparison with structural data and molecular dynamic simulations. HEK cells are an ideal system for this as they are easily cultured, transfected and amenable to patch clamp. However, HEK cells express endogenous human ASIC1 which may complicate interpretation. Therefore, we removed the endogenous human ASIC1 using CRISPR. To do this, exon 2 of the human ASIC1 gene was targeted with a guide-RNA cloned into a Cas9-GFP expressing vector (see Methods). Single GFP-positive HEK cells were clonally expanded and screened using PCR followed by sequencing (Supplemental Figure 1A). One such clonal population was selected for further characterization. As seen in Supplemental Figure 1, this cell line had negligible ASIC1 immunoreactivity compared to either wild type HEK cells or HEK cells transfected with human ASIC1a. Furthermore, whole cell patch clamp recordings from these presumptive KO cells found no significant currents in response to pH 5 application. All other experiments in this study used this HEK ASIC1 KO cell line where endogenous ASIC1 has been removed.

To investigate the kinetic consequences of Q277G in cASIC1, we excised outside-out patches from HEK KO cells transfected with either wild type cASIC1 or cASIC1 Q277G, along with eGFP. Patches were jumped from pH 8, to populate the resting state, into pH 6.5, 6 or 5.5, to activate and desensitize the channels. We were surprised to find that desensitization is completely intact in Q277G (Figure 1C). Indeed, the rate of desensitization was accelerated more than two-fold (Figure 1C). We also noted that there was a slight elevation of the steady-state or equilibrium current with pH 5.5 stimuli (Figure 1C and F, %steady-state current: wild type 0.09 ± 0.03%, n = 6; Q277G 2.0 ± 0.6%, n = 7, p = 0.005). To better compare with past work^24^, we used the same pH 6.5 stimulus. Interestingly, the elevated steady-state current was more prominent with less acidic stimuli, increasing to approximately 10% of the peak response using pH 6.5 (Figure 1D-F, %steady-state current: pH 6.0 1.8 ± 0.4%, pH 6.5 10 ± 2%, n = 7, p = 0.0001). Such a pH-dependent increase in steady-state current was not detectable in wild type channels, although the amplitudes of these steady-state currents are exceedingly small and hence difficult to measure (Figure 1D, F, %steady-state current: pH 6.0 0.15 ± 0.09%, pH 6.5 0.06 ± 0.04%, n = 6, p = 0.32 versus pH 5.5 steady-state).

The robust desensitization of Q277G was unexpected given prior work. However, we did observe a small yet significant increase in the current at steady-state, particularly at more alkaline stimulating values (Figure 1E-F). We hypothesize that this phenotype arises from a weaker pH-dependence of recovery from desensitization. ASIC recovery from desensitization is strongly dependent on the pH between the conditioning and test stimuli. Relatively alkaline inter-stimuli pH values accelerate recovery while more acidic inter-stimuli pH values slow recovery^23,47–49^. If one extrapolates this trend, then at more acidic values (i.e., pH 5.5) recovery is very slow and transitions from the desensitized state to the open or resting states are very unfavorable. Consequently, there is minimal steady-state current. The elevated steady-state current of Q277G suggests that Q277G recovery may be faster than wild type and/or less influenced by inter-stimuli pH values. To test this, we examined Q277G recovery from desensitization using several inter-stimuli pH values in the same patch. Consistent with our hypothesis, Q277G recovery from desensitization is substantially faster than wild type cASIC1 (Figure 2). Specifically, Q277G recovery had a time constant of 2.03 ± 0.05 ms (n = 5) at pH 8 which is roughly 400 fold faster than wild type cASIC1 (840 ± 90 ms, n = 5, p < 1e^−5^)^23^. Furthermore, the recovery time constants remained very fast at pH 7.4 and 7.0 (τ_rec_(pH 7.4) = 4.7 ± 0.2 ms, τ_rec_(pH 7.0) 34 ± 2 ms, n = 5). Thus, these data support the notion that the elevated steady-state current in Q277G arises from faster recovery from desensitization in general.

**Figure 2.**
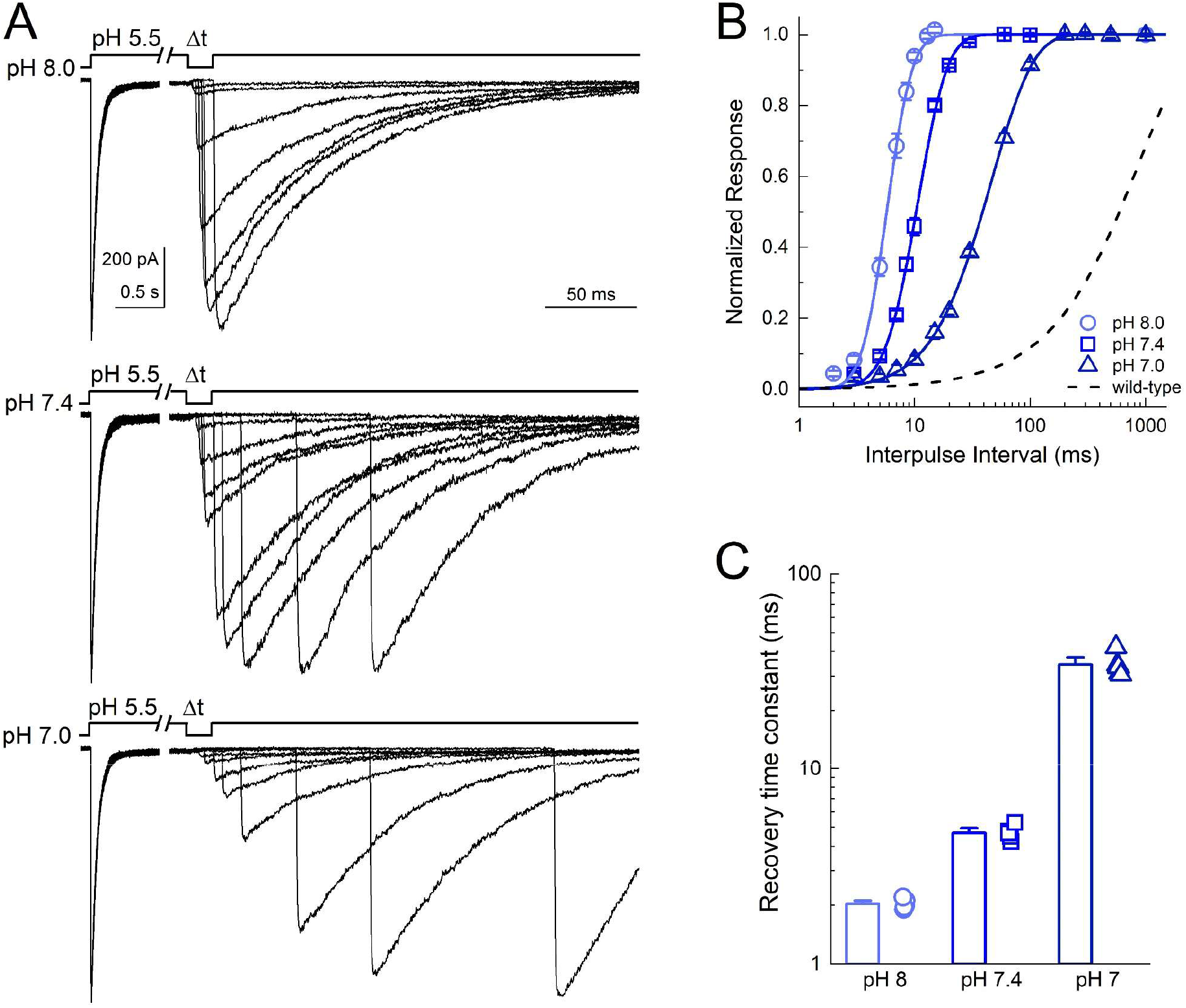
Q277G rapidly recovers from desensitization over a wide pH range. **(A)** Outside-out patch recordings of cASIC1 Q277G recovery from desensitization with interpulse pH values of 8.0, 7.4 and 7.0 (*upper, middle and lower traces, respectively*). All data from the same patch. Note the break and change in time base between conditioning and test pulses. **(B, C)** Summary recovery curves **(*B*)** and time constants **(*C*)** for Q277G recovery at different interpulse pH values. All pH values tested in the same patch. Symbols denote individual patches and error bars show S.E.M. Dotted line is the recovery time course of wild type cASIC1 with pH 8 drawn from Rook et al., 2020a.

Since steady-state desensitization (SSD) at any given pH value reflects a balance between channels entering and exiting the desensitized state, we hypothesized that the four-hundred fold faster recovery from desensitization would lead to a notable right-shift in the SSD curve. To test this, we constructed both activation and inhibition curves of Q277G and wild type cASIC1 in excised patches (Figure 3). We found that the pH-dependence of activation of Q277G was slightly more alkaline compared to wild type (wild type pH_50_ = 6.51 ± 0.01, n = 5; Q277G pH_50_ = 6.55 ± 0.01, n = 6, p = 0.006, Figure 3). However, the steady-state desensitization of Q277G was considerably right-shifted (Figure 3B-D). Specifically, the pH_50_ of SSD shifted from 7.30 ± 0.01 in wild type to 6.70 ± 0.01 in Q277G (n = 6 for both, p < 1e^−5^). The magnitude of the right-shift was sufficiently large to induce overlap with the activation curve. This distinct ‘window current’ led to standing currents with ‘baseline’ pH values such as pH 6.8 or 6.4 (Figure 3B) and a pronounced ‘foot’ on the acidic side of the SSD curve (Figure 3C). Taken together, we have found that Q277G produces only a small reduction in desensitization (or enhanced steady-state current). Further, that Q277G dramatically accelerates recovery from desensitization and right-shifts SSD curves with minimal effect on activation curves. We hypothesize that the recovery and SSD phenotypes all result from reducing the stability of the desensitized state. It has previously been suggested that the conformation of Gln276 (human ASIC numbering) controls the stability of the desensitized state by acting as a valve or steric barrier to regulate isomerization of the β11-12 linker^24^. To gain insight into the structural mechanism, we turned to molecular dynamic simulations.

**Figure 3.**
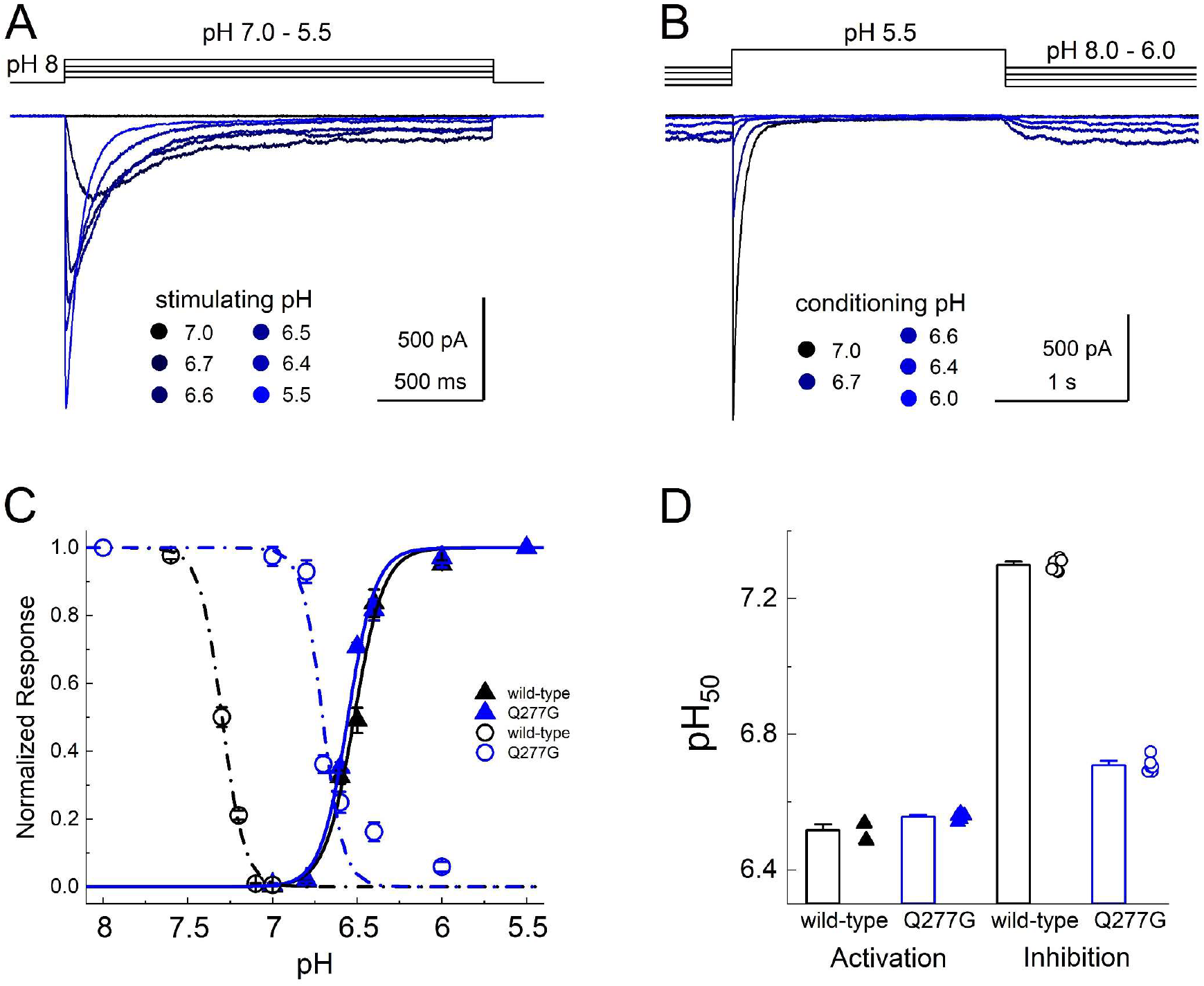
Q277G right-shifts steady-state desensitization without altering activation. **(A)** Outside-out patch recording of cASIC1 Q277G responses to increasingly acidic solutions. Darker solutions are more basic while acidic solutions are more blue. **(B)** Responses of Q277G to pH 5.5 application when preincubated with solutions ranging from pH 8 to 6. Note that solutions of intermediate acidity (pH 6.8-6.4) produces persistent currents at equilibrium. **(C)** Response curves to activation (*solid triangles*) or steady-state desensitization (*open circles*) for wild type (*black*) or Q277G (*blue*). **(D)** Mean ± SEM pH_50_s of activation (*left*) and steady-state desensitization (*right*) for wild type (*black*) or Q277G (*blue*). Fits from individual patches are shown as symbols following the legend **(C).**

Examining the proposed desensitized state structure of cASIC1 suggested that the Gln277 sidechain might form a hydrogen bond to the backbone oxygen atom of Leu414 when the linker is in the desensitized conformation (Figure 4A). Rather than acting as a valve, Gln277 could potentially stabilize the desensitized conformation through this hydrogen bond as we proposed from previous simulations^23^. Additionally, three acidic residues, Glu80 and Glu417 in the lower palm domain and Glu412 in the upper palm domain, are within potential hydrogen bond distance of Gln277, partly depending on the protonation states of the acidic residues. Thus, the structure suggests that Glu277 could play a role in a larger hydrogen bond network (Figure 4A). Because protonation states of the acidic side chains cannot be observed but must be inferred, we first tested the relative stability of the potential hydrogen bonds in the presence of different protonation states of Glu80, Glu412 and Glu417. Each residue can be protonated or not, giving rise to eight possible protonation combinations. Using the desensitized state structure (PDB: 4NYK)^27^ as a starting point, we simulated each protonation scheme for three repeats of 100 ns each. To quantify the stability of potential hydrogen bond interactions between the residues of interest, we measured the fraction of time that each potential hydrogen bond was present over the course of the simulations. Potential hydrogen bond donors considered were the side chains of Glu80, Gln277, Glu412 and Glu417, while potential hydrogen bond acceptors were the same side chains along with the backbone oxygen atom of Leu414. An interaction was considered as a hydrogen bond when the donor-acceptor distance was within 3.0 Å and the donor-hydrogen-acceptor angle greater than 150°. The overall hydrogen bond analysis (Supplemental Figure 2) illustrated that no matter the protonation states, Gln277 very rarely acted as a hydrogen bond acceptor. On the contrary, Gln277 often participated as a hydrogen bond donor in fairly stable hydrogen bonds. Looking at all 72 chains analyzed (8 setups × 3 repeats × 3 chains), Gln277 formed hydrogen bonds of varying stability to Glu80 in 75% of the cases; to Glu412 in 25% of cases; to L414 in 90% of cases and to Glu417 in 35% of cases. Therefore, we deemed the hydrogen bonds to Glu80 and to Leu414 to be most important. These two hydrogen bonds showed the highest stability in the setup in which Glu412 and Glu417 were protonated while Glu80 was deprotonated (E80-/E412H/E417H in Supplemental Figure 2). Thus, we chose this protonation setup to be the most stable for the desensitized state. Under these conditions, the side chain conformation of Gln277 was generally stable and positioned to hydrogen bond with the side chain of Glu80 and the backbone carbonyl oxygen of Leu414 (Figure 4B, Supplemental Movie 1). These interactions are noteworthy as mutations of either Glu80 or Leu414 can profoundly alter desensitization kinetics^23,50–53^. In particular, motion of Leu414 is a critical regulator of ASIC desensitization, underscoring the potential significance of these contacts. Figure 4 and Supplemental Figure 3 illustrates this analysis, showing that Q277 spends considerable time in putative hydrogen bond interactions with both Glu80 and Leu414. We hypothesized that such a network stabilizes the desensitized state with Gln277 acting as a critical hub. This role of Gln277 as an electrostatic hub is in contrast with the purely steric ‘valve’ model of Gln277 proposed previously^24^. We reasoned that a Q277N mutation may delineate between these hypotheses. If the ‘steric’ hypothesis is true, then shortening the side chain (Q277N) should produce minimal effect on desensitization kinetics. However, if the electrostatic hub model is more accurate, then the sub-optimal bonding distances of Q277N should result in much faster recovery from desensitization. To confirm that Q277N does attenuate hydrogen bond interactions, we repeated simulations using the Q277N mutation and observed that Q277N showed a greatly reduced capacity to participate in hydrogen bonds with Glu80 and Leu414 (Figure 4C-F, Supplemental Movie 2, Supplemental Figure 3). Therefore, we measured the recovery from desensitization of Q277N in excised patches. Consistent with the electrostatic hub hypothesis, Q277N recovers from desensitization much faster than wild type at all pH values tested. Specifically, at pH 8 the recovery time constant for Q277N is 4.0 ± 0.1 ms (n = 7, p < 1e^−5^ versus wild type, Figure 5). This is slowed to 32 ± 3 ms and 1500 ± 150 ms at pH 7.4 and 7 (n = 6 and 5, respectively; Figure 5). Next, we eliminated any residual capacity of the 277 position to participate in hydrogen bonds by using the Q277L mutation, which has identical steric factors as Q277N but no capacity for electrostatic interactions with nearby side chains. Consistent with the electrostatic hub hypothesis, Q277L has comparable recovery kinetics to Q277N (τ_rec_(pH 8) = 3.5 ± 0.1 ms, τ_rec_(pH 7.4) 39 ± 2 ms, τ_rec_(pH 7) = 2400 ± 90 ms n = 5-7, Figure 5B-D). While these time constants are slower than Q277G, they are orders of magnitude faster than wild type, suggesting the essential feature of Gln277’s function is as a hydrogen bond hub or coordinator and not a steric valve.

**Figure 4.**
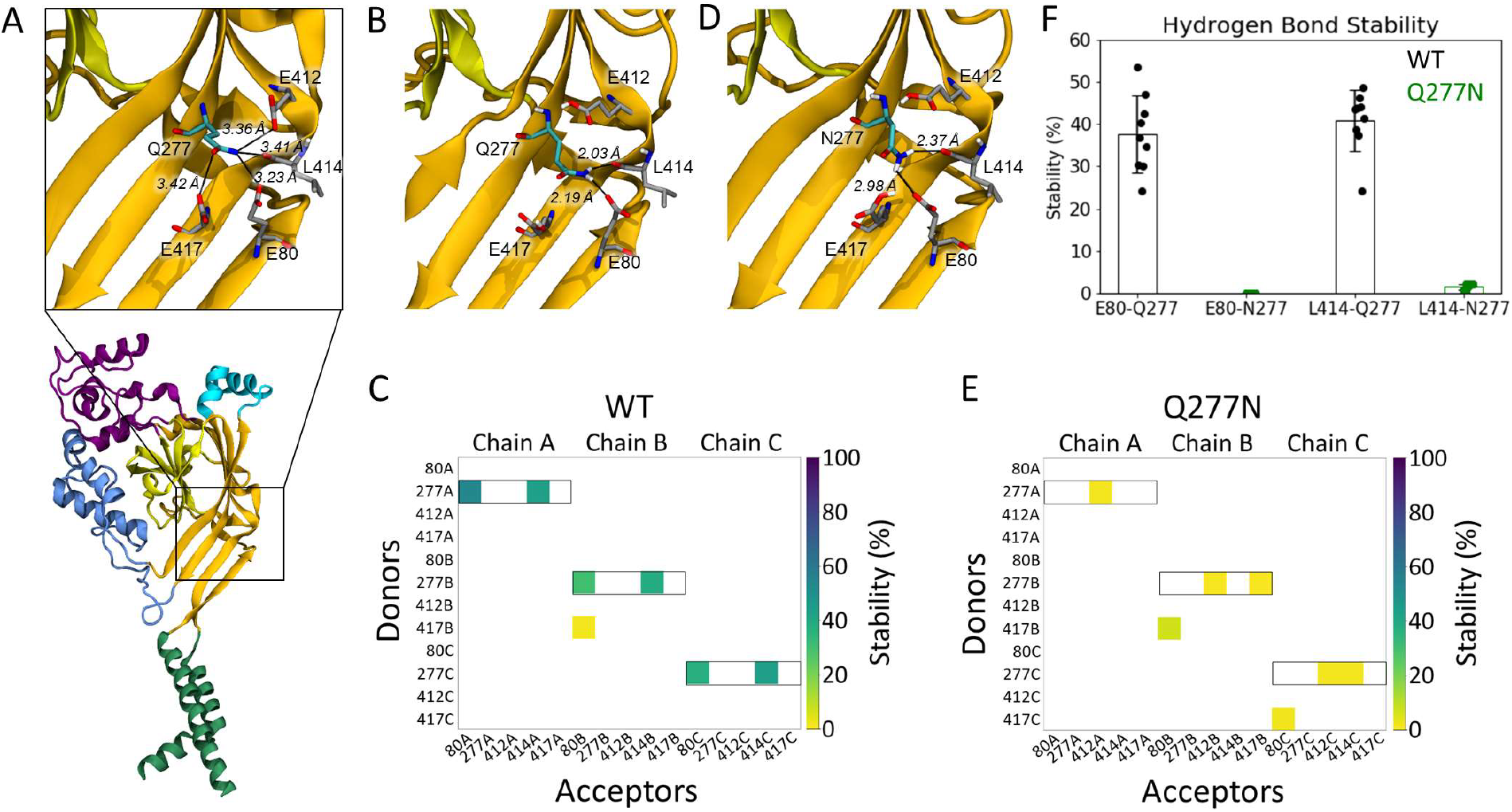
Gln277 links Leu414 and Glu80 via hydrogen bond network. **(A)** A single subunit of cASIC1 in the desensitized state, illustrating residues within potential hydrogen bonding distance of Gln277 (*inset*). Colors as in Figure 1A. **(B)** Snapshot from a WT simulation illustrating the hydrogen bond network with Gln277 in the center, hydrogen bonding to L414 and E80. The snapshot was taken at 8.6 ns. **(C)** Hydrogen bond analysis for a representative repeat (100 ns) of wild type with E80 deprotonated and E412 and E417 protonated. All hydrogen bonds formed between donors and acceptors of the sidechains of E80, Q277, E412 and E417 are considered, as well as hydrogen bonds in which the backbone oxygen atom of L414 participates as an acceptor. Acceptors are listed horizontally, donors vertically. The colored squares illustrate that a given hydrogen bond is present for part of the 100 ns of simulation, following the color bar given to the right. Hydrogen bonds in which Q277 participates as a donor are highlighted by black boxes. **(D)** Snapshot from a Q277N simulation illustrating that the inserted Asn residue is too short to form the same hydrogen bond network as Gln277. The snapshot was taken at 19.2 ns. **(E)** Hydrogen bond analysis as **D**, but for the Q277N mutant. **(F)** Average stability (*bars*) of the E80-Q277 and the L414-Q277 hydrogen bonds in the wild type (*black*) and Q277N simulations (*green*) (WT: E80-Q277: 38 +/− 9%; L414-Q277: 41 +/− 7%; Q277N: E80-N277: 0.02 +/0.01%; L414-N277: 1.6 +/− 0.6%. The nine data points (3 chains × 3 repeats) are illustrated as points and the error bar depict standard deviation.

**Figure 5.**
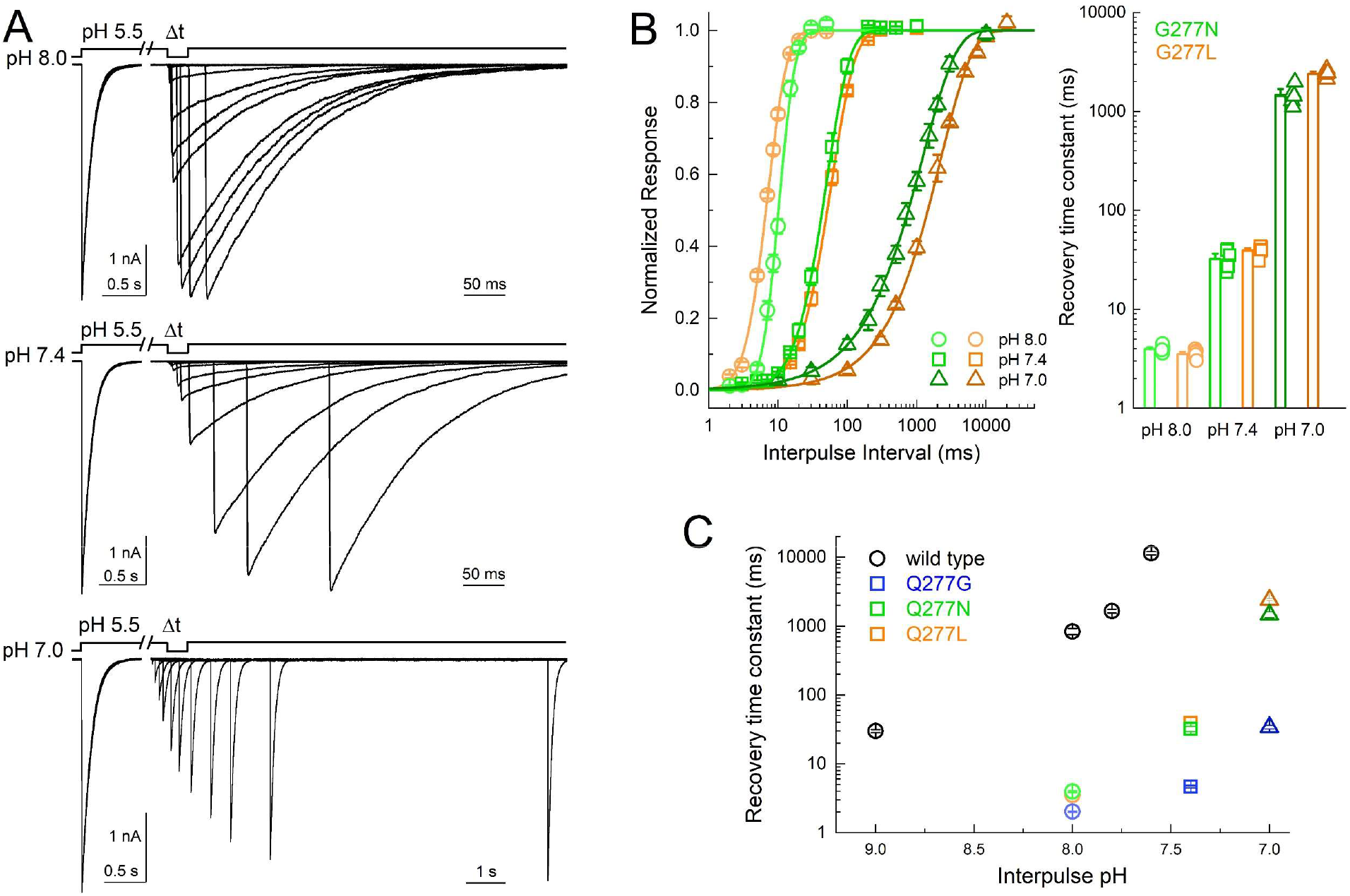
Q277N recovers nearly as fast as Q277G. **(A)** Outside-out patch recordings of cASIC1 Q277N recovery from desensitization with interpulse pH values of 8.0, 7.4 and 7.0 (*upper, middle and lower traces, respectively*). All data from the same patch. Note the break and change in time base between conditioning and test pulses. **(B)** Summary recovery curves (*left*) and time constants (*right*) for Q277N recovery at different interpulse pH values. All pH’s tested in the same patch. Symbols denote individual patches and error bars show S.E.M. **(C)** Summary of recovery time constants at various pH values for wild type, Q277G and Q277N. Wild type data drawn from Rook et al., 2020a.

These data demonstrate that in cASIC1 Q277G does not block desensitization. Rather, Q277G induces a slight increase in steady-state current that is pH-dependent. Given this, we re-examined the Q276G mutation in human ASIC1a as was previously published, as well as mouse ASIC1a. In both cases, the Q276G mutation gave small, barely detectable currents in excised patches, necessitating whole cell recording. In the case of mouse ASIC1a Q276G, even whole cell currents were too small to resolve and examine convincingly (43 ± 13 pA, n = 10). Therefore, we confined ourselves to hASIC1a Q276G. As with cASIC1 Q277G, desensitization was intact in this mutant but, rather than accelerating current decay as in cASIC1, hASIC1a Q276G showed much slower decay kinetics with pH 5.5 evoked responses (Figure 6). Moreover, currents evoked by pH 6.5 did not exhibit macroscopic desensitization on these time scales. Interestingly, we observed rather fast rundown or inhibition when using a 5 second stimulus and 20 second intervals (Figure 6A). This stimulus and interval duration has proven adequate for wild type hASIC1a in our hands. To properly measure the desensitization time course and allow for complete recovery, we progressively extended both the stimulus and interval times. Ultimately, using a 100 second pH application spaced by 120 seconds, we found that hASIC1a Q276G channels desensitize very strongly using pH 5.5 or less strongly when using pH 6.5 (Figure 6, steady-state current: pH 5.0 3.7 ± 0.5% of peak; pH 6.5 17 ± 2% of peak, n = 5). However, their desensitization time course is considerably longer than wild type (Q276G: 8800 ± 1400 ms, n = 5; wt: 788 ± 104 ms, n = 5, p < 1e^−5^). Taken together, we demonstrate that the Q/G mutation does not abolish desensitization as previously reported. Rather, in cASIC1 this mutation elevates the steady-state current, accelerates recovery from desensitization and reduces the stability of the desensitized state. Molecular dynamic simulations and subsequent mutagenesis suggest these phenotypes arise by destabilizing a critical hydrogen bond network, which in the wild type stabilizes the desensitized state. In hASIC1a, this mutation also does not abolish desensitization yet the functional phenotype is distinct from cASIC1.

**Figure 6.**
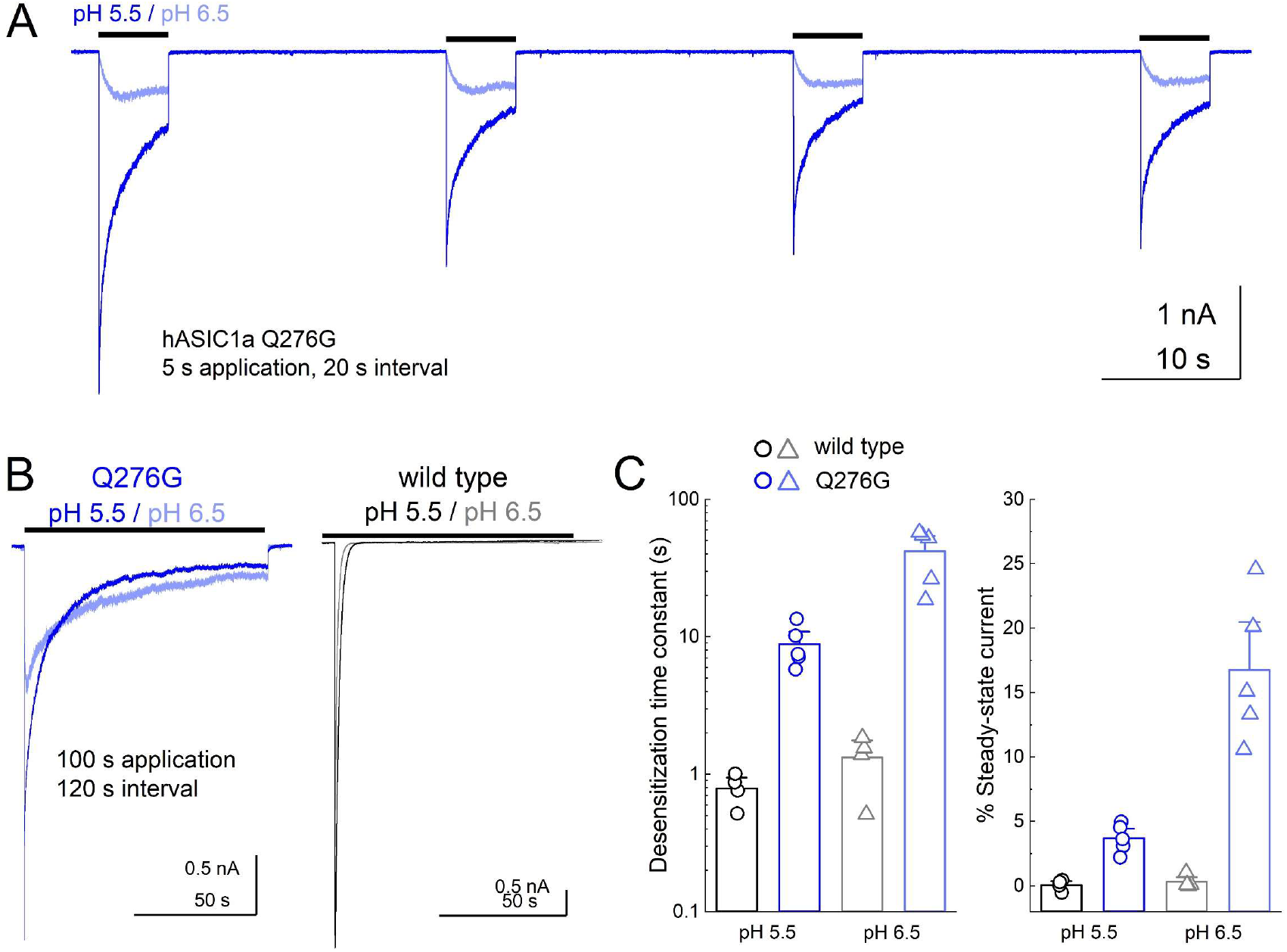
Human ASIC1a Q276G also does not abolish desensitization. **(A)** Whole cell recording of hASIC1a Q276G during repeated applications of pH 5.5 (*blue trace*) or 6.5 (*light blue trace*). **(B)** Q276G (*left, blue traces*) and wild type (*right, black traces*) responses to longer pH 5.5 and 6.5 applications with greater intervals. **(C)** Summary of desensitization time constants (*left*) and percent of steady-state current (*right*) at pH 5.5 (*circles*) and 6.5 (*triangles*) for wild type (*black*) and Q277G (*blue*). Symbols denote single cells and error bars are SEM.

## Discussion

We explored the properties of the recently described Q276G ASIC1 mutation^24^ (human numbering) using a combination of fast perfusion electrophysiology and molecular dynamics simulations. In contrast to prior work, we find that this mutation does not abolish ASIC1 desensitization. Rather, this mutation leads to a slight elevation in steady-state current that is more pronounced with weaker pH stimuli (Figure 1). In cASIC1, Q277G also markedly accelerates recovery from desensitization over a wide pH range (Figure 2) and right-shifts the pH-dependence of steady-state desensitization without substantially altering activation (Figure 3). All-atom simulations of the cASIC1 desensitized state indicate that this conformation is stabilized by a network of hydrogen bonds linking the lower palm residue Glu80, through Gln277, with the β11-12 linker (Figure 5). Consistent with this, compromising the hydrogen bond network by shortening the Q277 side chain either slightly (Q277N) or significantly (Q277G) has a profound impact on the stability of the desensitized state as measured by recovery from desensitization (Figure 5). Finally, we found that hASIC1a Q276G also desensitizes but both enters and exits the desensitized state slower than wild type hASIC1a (Figure 6).

### Comparison with previous studies

The original report that Q276G blocks desensitization used human ASIC1a in a *Xenopus* oocyte expression system primarily using bath perfusion, pH 6.5 as a stimulus with pH 7.4 as a baseline pH^24^. Using pH 6.5 as a stimulus, combined with the phenotype of hASIC1a Q276G, may have led to the assertion that Q276G blocks desensitization. Specifically, the slow desensitization of hASIC1a Q276G and elevated steady-state current produced by pH 6.5 can lead to an observed lack of desensitization or current decay during shorter agonist applications (Figure 6A). This problem may be exacerbated by the slow recovery of hASIC1a Q276G, particularly when using pH 7.4 as a baseline pH, thus leading to the suppression or lack of recovery of the peak while allowing the steady-state to persist. We suggest that this experimental setup, combined with the phenotype of hASIC1a Q276G, led to the conclusion that desensitization was abolished.

Rather than Gln277 controlling desensitization and recovery by the proposed valve mechanism^24^, we provide evidence that Gln277 is central to an important hydrogen bond network linking the influential Glu80 residue in the lower palm with the critical β11-12 linker that governs desensitization. How might such a network function in the ASIC gating cycle? In our simulations both Glu412 and Glu417 are protonated, leaving the Gln277 amide to act as a hydrogen bond donor to the deprotonated Glu80 and the backbone carbonyl of Leu414. The interaction with the carbonyl is the most commonly observed (Figure 4, Supplemental Figures 2 and 3). We propose that in the desensitized state, Gln277 partly contributes to the stability of Leu414 by this hydrogen bond, with Gln277 being held in this advantageous position by Glu80. Upon alkalization either Glu412, Glu417 or both tend to become deprotonated, acting as alternative hydrogen bond acceptors and thereby helping to pull the amide group of Q277 away from the backbone carbonyl of Leu414, releasing Leu414. This would facilitate recovery from desensitization. However, it is difficult to reconcile this hypothesis with the hASIC1a Q276G data which shows an apparent slowing of recovery from desensitization.

### Human versus chicken data

In our hands, the Q/G mutant gives opposite effects in cASIC1 versus hASIC1a, accelerating kinetics in the former but slowing them in the latter (Figure 1 versus Figure 6). This is reminiscent of the effects of psalmotoxin which inhibits mammalian ASICs by stabilizing a desensitized state^54^ yet activates cASIC1, promoting an unusual non-selective open state^55,56^. Another recent example is the blunted effect of mambalgin in cASIC1 compared to hASIC1a, which can largely be reversed by several point mutations^57^. Presently it is unclear what the source of these differences is. Human and chicken ASIC1 contain 56 amino acid differences, 31 of which are in the extracellular domain. A number of these are concentrated in the wrist region, including a 2-amino acid insertion. Given the wrist region’s involvement in gating^58^, it is possible that many species-specific differences arise from here. Further differences relevant for our kinetic experiments include the TRL versus SQL substitutions around amino acids 84 to 86^59^ as well as Ser275Ala, Val368Leu and Ala413Val which are all relatively proximal to Gln277 (chicken to human differences). We hypothesize one or more of these changes subtly alters the structure of hASIC1a, potentially imparting distinct pK_a_ values on critical palm residues and thus changing the phenotype of Q276G. As more hASIC1a structures become available in distinct functional states^57^, we hope to explore the source of these differences and the conservation of mechanisms in more detail. A similar examination may uncover why the equivalent Q269G mutation in ASIC3 does appear to inhibit desensitization even with pH 5^60^. Regardless of phenotypic differences, our data clearly indicate that both cASIC1 and hASIC1a Q277G mutants desensitize to a large extent. Therefore, using these mutations to explore either biophysical mechanisms of desensitization, or its physiological consequences, may be problematic.

## Acknowledgements

Funding for this work was provided by NIH T32GM068411-15 and Joan Wright Goodman Fellowship to M.L.R., NSF GRFP to T.C., NSERC Discovery Grant (RGPIN 2019-06864) and Canada Research Chairs grant (950-232154) to M.Mu and NIH R00NS094761, R35GM137951 and NARSAD Young Investigator Award to D.M.M. We thank Dr. Matthieu Chavent for initial discussions concerning the hydrogen bond analysis.

## Author contributions

M.L.R., M.Mi., D.K., T.C. and D.M.M. conducted experiments and analyzed data. M.L.R., M.Mi., M.Mu. and D.M.M. interpreted results and edited the manuscript.

---

**Supplemental Figure 1.**
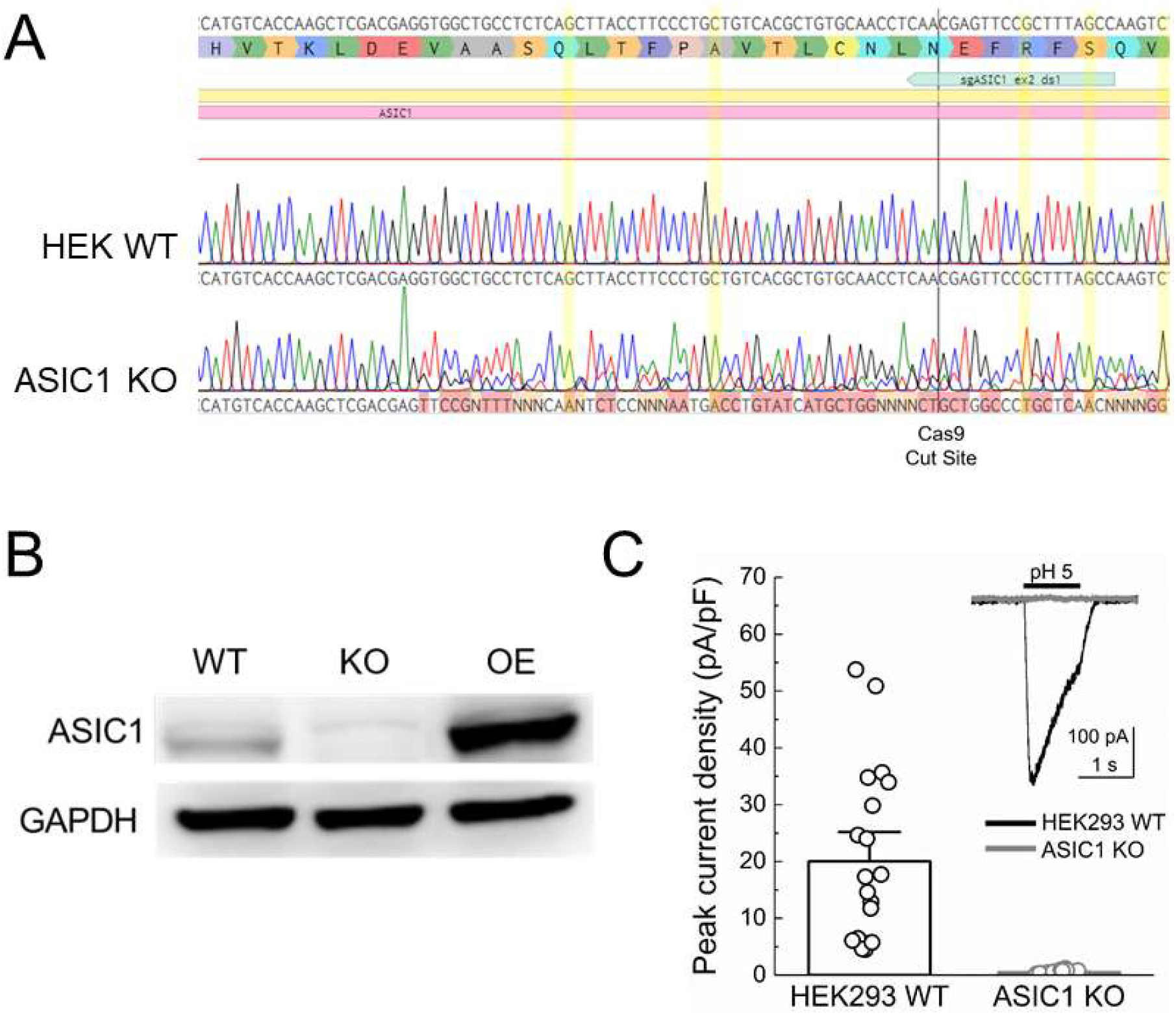
Validation of human ASIC1 knockout cells. **(A)** Sequencing of genomic DNA from HEK wild type cells (WT) or ASIC1 knockout cells (KO) of human ASIC1 gene’s second exon. Positions highlighted in yellow show that all alleles of the knockout line have been frame shifted. **(B)** Western blot of wild type HEK293 cells (WT), ASIC1 knockout cells (KO) and KO cells overexpressing human ASIC1a (OE). **(C)** pH-5 evoked whole cell current densities from HEK wild type (*black*) and KO (*grey*) cells. Raw traces are inset. pH 5-evoked peak current density: wild type 20 ± 3 pA/pF, n = 20 cells; KO pH 5 0.51 ± 0.05 pA/pF, n = 20 cells; p < 1e^−5^, Mann-Whitney U test. Circles represent individual cells and error bars depict S.E.M.

**Supplemental Figure 2.**
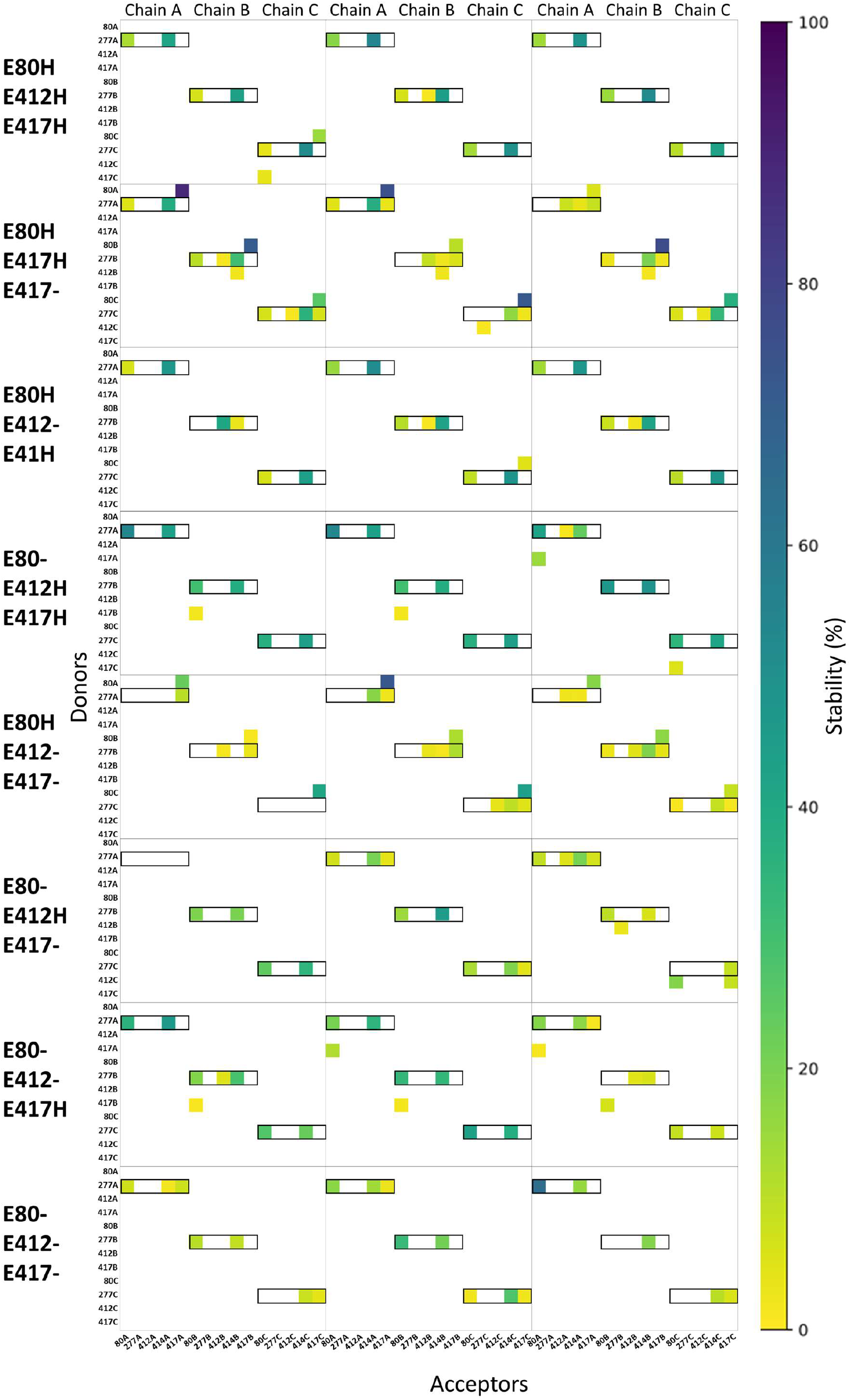
Hydrogen bond analysis for all eight possible protonation setups concerning E80 (H/-), E412 (H/-) and E417 (H/-). All hydrogen bonds formed between donors and acceptors of the sidechains of E80, Q277, E412 and E417 are considered, as well as hydrogen bonds in which the backbone oxygen atom of L414 participates as an acceptor. Hydrogen bond stability, following the color scale, is illustrated for each chain in each of three runs (100 ns in duration). Hydrogen bonds in which Q277 participates as a donor are highlighted by black boxes.

**Supplemental Figure 3.**
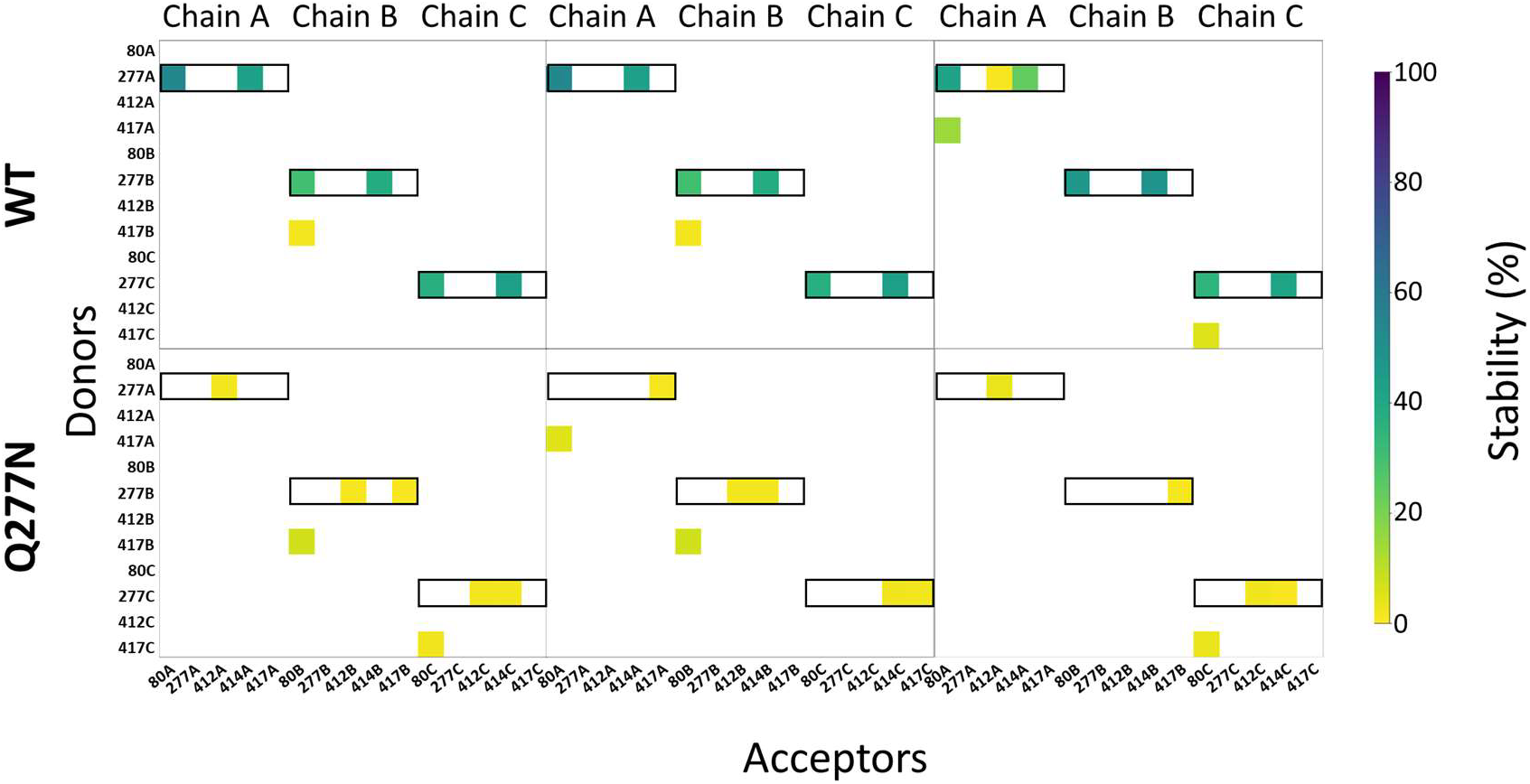
Q277N reduces hydrogen bond stability. Same analysis as Supplemental Figure 2. Hydrogen bond stability for each chain in each of three runs (100 ns in duration). Note that Q277N shows less stability than wild type of each chain and each run.

